# From acidic extreme environments to agriculture: a multi-stress-tolerant *Talaromyces pseudofuniculosus* enhances soybean growth under abiotic stress

**DOI:** 10.64898/2026.10.06.756979

**Authors:** Ivana Staiano, Stefany Castaldi, Marco Masi, Maria Michela Salvatore, Giorgio de Tommasis, Chiara Gentile, Anna Andolfi, Alessio Cimmino, Rachele Isticato

## Abstract

Extremophilic fungi are a relatively unexplored source of stress-resilient microbes for sustainable agriculture. Here, we describe *Talaromyces pseudofuniculosus* FAR, isolated from the highly acidic site of “Piccolo Inferno” in southern Italy and identified by ITS and β-tubulin (BenA) sequencing. Despite its origin from an extreme environment, *T. pseudofuniculosus* FRA remained fully functional under non-stress conditions and displayed tolerance to salinity and heavy metals while maintaining key plant growth-promoting traits, including phosphate solubilization, auxin-related compound production and hydrolytic activities. In soybean (Glycine max L.), the live fungus promoted plant growth under both non-stress conditions and agriculturally relevant salinity and copper stress, and successfully colonized the roots. A cell-free extracellular extract (exCFS) promoted early seedling development in vitro, but its effects were not sustained in pot experiments. GC–MS analysis provided a qualitative characterization of the extracellular metabolite profile and revealed condition-dependent differences between control, NaCl and CuSO₄ growth conditions, indicating environment-dependent modulation of fungal metabolism. Overall, these findings highlight the physiological and metabolic plasticity of *T. pseudofuniculosus* FRA and support its potential as a fungal biostimulant for saline- and metal-impacted agricultural soils.

## INTRODUCTION

The increasing incidence of soil degradation, salinization, and heavy-metal contamination is becoming a major bottleneck for sustainable agricultural productivity worldwide. In recent decades, agricultural soils have been progressively impacted by industrial emissions, intensive farming, irrigation with contaminated water, and long-term metal-based agrochemical use, leading to the accumulation of toxic metals such as cadmium (Cd) and copper (Cu) in high concentrations. Although Cu is naturally present in soils and plays an essential role in supporting plant growth, its excessive accumulation can become toxic to plants and interfere with nutrient uptake, reduce photosynthetic efficiency, and disrupt the cellular redox balance, thus reducing crop yield and quality (Burkhead *et al*., 2009). In parallel, salinity has emerged as one of the dominant abiotic stress factors strongly affecting millions of hectares of arable land, particularly in regions subject to irrigation, climate change, and soil degradation (FAO, 2015). High salinity leads to both osmotic stress, by limiting water extraction from the soil, and ionic stress, through the accumulation of toxic ions and metabolites that disrupt cellular metabolism and plant development (Munns and Tester, 2008). Current remediation strategies to alleviate these abiotic stresses, such as chemical amendments, soil replacement, or physical washing (Rahim *et al*., 2025), are often cost-intensive, environmentally invasive, and poorly scalable. These limitations have accelerated the demand for biologically based, low-input, and field-deployable solutions compatible with sustainable and regenerative agriculture frameworks. A sustainable strategy for alleviating heavy metals and salt stress in plants involves the use of microorganisms as biostimulants and biofertilizers (Staiano *et al*., 2025; Castaldi *et al*., 2023). This microbial biotechnological application is increasingly recognized as a high-TRL technology with strong commercial traction, supported by favourable regulatory frameworks (e.g. EU Regulation 2019/1009) and growing market demand (Vejan *et al*., 2016). Plant growth-promoting microorganisms (PGPM) can enhance crop performance under stress by improving nutrient bioavailability, modulating phytohormone balance, and reshaping rhizosphere chemistry. Certain PGPM can directly sequester or remove metals from contaminated environments, while concurrently synthesizing plant growth regulators, including gibberellins and auxins. This dual functionality enhances plant tolerance to abiotic and biotic stresses and contributes to improved plant growth and performance (Hassan *et al*., 2020). Other PGPM have been shown to produce osmoprotectants, which can alleviate salt stress and improve plant growth in saline environments (Khan, Hamayun, Kim, *et al*., 2011). Among candidate microorganisms that could promote plant growth in stressful environments, filamentous fungi have several strategic advantages for industrial development over bacteria, including robustness under fluctuating environmental conditions, high secretory capacity, and compatibility with solid-state or liquid fermentation processes (Patel *et al*., 2023). Their effectiveness is mainly attributed to their broad ecological distribution and ability to withstand environmental stresses (Deng and Cao, 2017). By occupying extreme habitats, such as acidic waters, metal-rich soils and saline niches, fungi have evolved mechanisms for physiological and genetic adaptation that confer cross-resistance to multiple stressors. These include enhanced antioxidant systems, altered cell-wall and membrane properties, specialized ion transporters, and metabolic pathways for metal detoxification or sequestration (Tiquia-Arashiro and Grube, 2019). In addition, their multicellular biomass allows efficient sequestration and chelation of metal contaminants, while the wide array of metabolites produced can enhance host plant growth in stressful environments (Khan *et al*., 2008; Hamayun *et al*., 2010). Consequently, fungi are emerging as ideal biostimulant-based platforms for biotechnological applications aimed at promoting plant growth and productivity while enhancing tolerance to metal stress and salinity. In this study, we isolated and characterized an extremophilic *T. pseudofuniculosus* strain from a highly acidic natural environment and evaluated its suitability as a microbial biostimulant under salinity and copper stress. As a plant model to assess the PGP effect of the strain, we used soybean (*Glycine max* L.), one of the world’s 16 major crops and a primary source of plant protein for human food and high-quality animal feed (Foley *et al*., 2011). It is estimated that the European Union produced 2.81 million metric tons of soybeans during 2023/2024 (United States Department of Agriculture), playing an essential role in both economic and dietary fields. Given its susceptibility to environmental fluctuations that significantly impair yield and quality, soybean represents a robust system for evaluating the plant growth–promoting and stress-mitigating potential of microbial inoculants (Bilal *et al*., 2019). By integrating stress-tolerance profiling, plant performance assays, and metabolomic analyses, we systematically evaluate both live-inoculum and cell-free (postbiotic) strategies, defining their translational potential for sustainable agriculture under environmental stress. Our work positions extremophilic fungi as realistic candidates for the development of robust, field-ready biostimulants targeting stressed and contaminated agricultural soils.

## Material and Methods

### Isolation and identification of fungal isolate

Samples of acidic water collected from “Piccolo Inferno” (Campania, Italy; 41.314487°N, 13.894006 °E; 29.5 °C; pH of 2.13) were kindly provided by Prof. Donato Giovannelli (Department of Biology, University of Naples “Federico II”, Italy). The samples were placed into sterile plastic bags and kept at 4 °C until processing. An enrichment step, adapted from selective approaches previously used for the recovery of fungi from highly acidic environments (Wang et al., 2021), was performed by incubating 1 mL of sample in 5 mL of acidified aqueous medium (pH 2.5, adjusted with 33% HCl) supplemented with 2% w/v of glucose at 25 °C, in order to selectively enrich acid-tolerant heterotrophic microorganisms. After 7 days of incubation, fungal biomass was observed and subsequently plated on Potato Dextrose Agar (PDA) (Difco, Fisher Scientific, Italy). Three replicate cultures of the isolate were incubated in the dark at 25 °C. Colony morphology (average colony diameter, colour, elevation, margin, shape, and surface) was evaluated after 7 and 14 days of incubation based on the morphotype concept (Kornerup and Wanscher, 1967; Bills, 1995; Lacap *et al*., 2003). Microscopy observations of the fungus were performed with an optical microscope (Olympus BH-2) equipped with a 40× objective. For molecular identification, DNA was extracted from fungal mycelium using the STAR protocol 1 (Conlon *et al*., 2022). Taxonomic identification was performed by sequencing both the Internal Transcribed Spacer (ITS) region and a partial sequence of the β-tubulin (BenA/Bt) gene. The ITS region was amplified using the universal primers ITS1 (5’-TCC GTA GGT GAA CCT GCG G-3’) and ITS4 (5’-TCC TCC GCT TAT TGA TAT GC-3’), whereas the β-tubulin gene was amplified using primers Bt2a (5′-GGTAACCAAATCGGTGCTGCTTTC-3′) and Bt2b (5′-ACCCTCAGTGTAGTGACCCTTGGC-3′) (Glass and Donaldson, 1995).

The amplified fragment was purified using the QIAquick PCR Purification Kit (Qiagen, Hilden, Germany) according to the manufacturer’s instructions and sequenced at the BMR Genomics sequencing facility (Padova, Italy). Sequence quality was analysed using SeqTrace; the minimum acceptable base call quality (Phred score) was set to 30, with a minimum of 20 consecutive bases at this threshold (Stucky, 2012). The trimmed and paired sequences were analysed using the Basic Local Alignment Search Tool (BLAST). The ITS and β-tubulin gene sequences were deposited in GenBank under accession number PX599580 and PX966871, respectively.

### Assessment of abiotic stress tolerance

To assess abiotic stress tolerance of the isolate, fungal plugs (5 mm in diameter) from one-week-old fungal colony grown on PDA plates were cut and inoculated in the centre of PDA plates containing increasing concentrations of CuSO_4_, PbCl_3_, NiSO_4_ (from 1 mM to 4 mM) and NaCl (from 0.1 M to 1.0 M), the media without the addition of metals or salt was used as a control. After 10 days of incubation (25 °C in darkness), the growth of the fungus was evaluated by measuring the diameter of the colony and by comparing it to the control. Results are expressed as Tolerance Index (percentage of growth of the fungus with respect to the control experiments). The experiment was conducted in triplicate.

### Evaluation of hydrolytic activities

Cellulose-, xylan-, and starch-degrading activities of the fungal isolate were evaluated using plate-based screening assays. Fungal plugs (5 mm in diameter) from one-week-old cultures grown on PDA plates were centrally inoculated onto basal media plates and incubated at 25 °C for 7 days. Enzymatic activity was assessed by the appearance of hydrolysis halos surrounding the colonies and quantified as the Enzymatic Index (EI) (Florencio *et al*., 2013, Flores *et al*., 2023), calculated as:

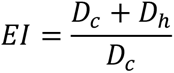

where *D_c_* is the colony diameter and *D*_ℎ_ is the diameter of the hydrolysis halo.

For cellulose and xylan assays, minimal media (MM composed of 1 g L^-1^ NaNO_3_, 1 g L^-1^ K_2_HPO_4_, 0.5 g L^-1^ MgSO_4_ · 7 H_2_O, 1 g L^-1^ KCl, 0.5 g L^-1^ Yeast Extract, 15 g L^-1^ agar) was supplemented with 0.5% (w/v) carboxymethyl cellulose sodium salt (Na-CMC) or 0.5% (w/v) beechwood xylan as the sole carbon source for cellulase or xylanase activity, respectively (Meddeb-Mouelhi *et al*., 2014). To evaluate enzymatic activity under abiotic stress conditions, both media were additionally supplemented with NaCl (0.1, 0.25, 0.5, and 1.0 M) or CuSO₄ (1.0, 2.0, 3.0, and 4.0 mM). After incubation, plates were flooded with aqueous Congo Red dye solutions (0.1% w/v for cellulase and 0.2% w/v for xylanase), incubated for 30 min in darkness, and then destained by washing twice with 1.0 M NaCl solution for 15 min (Maki *et al*., 2012). The appearance of yellow halos around the colonies indicated extracellular hydrolytic enzyme activity. For amylase activity, starch agar plates were used according to the literature (3 g L^-1^ beef extract, 10 g L^-1^ soluble starch, 15 g L^-1^ agar) as described by Lal and Cheeptham (2016). After incubation, plates were stained with iodine vapours, clear yellowish zone surrounding the colonies indicated starch hydrolysis. All the assays were conducted in triplicate.

### Quantification of glucanase activity

Endoglucanase and exoglucanase activities of the crude cell-free supernatant (CFS) were assayed according to Meng et al. (2021). The fungus was cultivated in 500 mL Erlenmeyer flasks containing 100 mL of PD broth at 25 °C, 120 rpm for 7 days. After incubation, the CFS was recovered by centrifugation (7000 g for 30 min at 4 °C) and sequential filtration through cotton and filter paper. Endoglucanase activity was determined in a reaction mixture containing 1.0 mL of CFS with 0.5 mL of 2.0% (w/v) solution of Na-CMC in 0.05 M acetate buffer (pH 4.5) and incubated for 30 min at 50°C (Jung *et al*., 2015). Exoglucanase activity was measured using a filter paper assay, in which a 1 × 6 cm strip of filter paper was incubated with 1.0 mL of CFS in 1.0 mL of 0.05 M acetate buffer (pH 4.5) at 50 °C for 1 h (De Castro *et al*., 2010; Ramani *et al*., 2012). Reducing sugars released after enzymatic hydrolysis were quantified using the 3,5-dinitrosalicylic acid (DNS) method (Miller, 1959; Meddeb-Mouelhi et al., 2014). Briefly, 3.0 mL of DNS reagent were added to each reaction mixture, followed by incubation at 100 °C for 5 min. Samples were then cooled, diluted with 20 mL of distilled water, and absorbance was measured at 540 nm. Acetate buffer was used as blank, and a glucose calibration curve (eight points ranging from 1.0 to 3.35 mg, in triplicate) was generated. One unit of endo- or exoglucanase activities was defined as the amount of enzyme catalyzing the release of 1 μmol of glucose per 1 min, under the assay conditions.

### *In vitro* evaluation of plant growth promoting properties

#### Determination of phosphate solubilization under salt and copper stress

Qualitative phosphate solubilization was checked by inoculating a 5-mm-diameter fungal plug of *T. pseudofuniculosus* FRA on Pikovskaya plates (Pikovskaya, 1948) adjusted to pH 7.0 (10.0 g L^-1^ glucose, 5.0 g L^-1^ Ca_3_(PO_4_)_2_, 0.5 g L^-1^ (NH_4_)SO_4_, 0.2 g L^-1^ NaCl, 0.1 g L^-1^ MgSO_4_ · 7 H_2_O, 0.2 g L^-1^ KCl, 0.002 g L^-1^ MnSO_4_ · 7 H_2_O, 0.002 g L^-1^ FeSO_4_ · 7 H_2_O, 0.5 g L^-1^ yeast extract, 10 mg L^-1^ bromophenol blue, 15 g L^-1^ agar). Plates were supplemented with NaCl (0.1 M, 0.5 M and 1.0 M) or CuSO_4_ (1 mM, 2 mM, 3 mM and 4 mM) and incubated at 25 °C in darkness for 6 days. Pikovskaya plates (PVK) without salt or metal supplementation were used as controls to evaluate phosphate solubilization under non-stress conditions. Phosphate solubilization was quantified by calculating the Phosphate Solubilization Index (PSI) according to Edi Premono et al. (1996), using the following formula:

Solubilization index = (colony + halo)_diameter_ / colony_diameter_

#### Production of Indole-3-Acetic Acid (IAA) and IAA-Related Compounds under salt and copper stress

IAA production and IAA-related compounds were determined under *in vitro* conditions. Erlenmeyer flasks (250 mL) containing 50 mL of Tryptophan Liquid Medium (TLM: 10.0 g L^-1^ casein hydrolysate, 5.0 g L^-1^ NaCl, and 1.0 g L^-1^ L-tryptophan, Sigma-Aldrich) were inoculated with 1.0 mL of 5-day-old PD broth culture of the fungal strain and incubated at 25 °C, 120 rpm, for 14 days. Other experiments were conducted with the same media (TLM) supplemented with NaCl (0.1 M and 0.25 M) or CuSO_4_ (1 mM and 3 mM); these concentrations were chosen because they did not interfere with the colorimetric assay. Uninoculated Erlenmeyer flasks were used as negative controls. Aliquots of the culture medium were collected after 7 and 14 days of growth to determine the concentration of IAA and IAA-related compounds. For quantification, the culture medium was filtered through a 0.22 μm syringe. Then, 2.0 mL of the filtrate were mixed with 4.0 mL of Salkowski’s reagent (H_2_O:H_2_SO_4_:FeCl_3_ 0.5 M in a ratio 50:30:1 respectively) (Gordon and Weber, 1951). The mixture was incubated for 30 min at room temperature, and colour development was measured by reading the absorbance at 530 nm. IAA concentration was determined by comparison with an eight-point calibration curve prepared using a methanolic IAA standard ranging from 1 to 100 μg mL⁻¹ (Sigma-Aldrich).

### Extraction of organic compounds

Cell-free supernatants (CFS) were obtained after 7 days of growth in 1L PD broth (control condition) and supplemented with 0.1 M NaCl (salt stress condition) or 1mM CuSO₄ (heavy-metal stress condition). CFS samples were sequentially filtered through cotton and filter paper, followed by filtration through 0.22-μm pore-size membranes (Corning®, New York, NY, USA), and divided into 50-mL aliquots. Selected aliquots were used for the extraction of extracellular organic compounds. The pH of each CFS was adjusted to 7 with 0.1 N NH₄OH, and each aliquot was extracted three times with an equal volume (50 mL) of ethyl acetate. The organic phases were combined, dried over anhydrous Na₂SO₄, and evaporated under reduced pressure to obtain the corresponding CFS extract (exCFS).

### Identification of secondary metabolites through GC-MS analysis

The crude extracts were analysed by GC-MS after trimethylsilylation with *N*, *O*-bis(trimethylsilyl)trifluoroacetamide (BSTFA) (Fluka, Buchs, Switzerland) as previously reported by Salvatore *et al*. (2024). In particular, 100 µL of BSTFA and 100 µL of acetonitrile were added to the organic extract, which was derivatized by incubation for in a microwave oven for 2 min at 180 W. GC-MS analyses were conducted with an Agilent 6850 gas chromatograph (Milan, Italy), equipped with an HP-5MS capillary column (stationary phase: 5%-phenyl-95%methylpolysiloxane; 30 m Length; 0.25 mm i.d.; 0.25 µm Film Thickness), coupled to an Agilent 5973 Inert mass selective detector operated in the full-scan mode (*m/z* 40–550) at a frequency of 3.9 Hz he electron ionization (EI) source and quadrupole temperatures were set at 200 °C and 250 °C, respectively. Helium was used as the carrier gas at a constant flow rate of 1.0 mL min⁻¹. The injector temperature was set at 250 °C. The oven temperature program was as follows: 70 °C for 1 min; ramped at 10 °C min⁻¹ to 170°C; then increased at 30 °C min⁻¹ to 280 °C and held for 5 min. The solvent delay was 4 min. Metabolites were identified by comparing their EI mass spectra at 70 eV with mass spectra collected in the NIST 20 mass spectral library (available at https://www.nist.gov/srd/nist-standard-reference-database-1a; accessed on 14 September 2025). Moreover, the identification was supported by the Kovats retention index (RI) calculated for each analyte by the Kovats equation, using the standard *n*-alkane mixture in the range C7-C40.

### Plant experiments

In all the experiments with *Glycine max* L., soybean seeds were sterilized as described in (Barbulova *et al*. (2005). After sowing on 0.5x Murashige and Skoog (MS) medium, unsynchronized seedlings were discarded three days after germination, and the remaining seedlings were maintained under axenic conditions at 25 °C in darkness (Murashige and Skoog, 1962).

### *In vitro* germination assay on *Glycine max* L. under physiological and abiotic stress conditions

The effects of *T. pseudofuniculosus* FRA and its exCFS on soybean (*Glycine max* L., Merr., cv. Sultana) seed germination were evaluated using complementary in vitro assays under both physiological and abiotic stress conditions. In all experiments, sterile seeds were grown on 0.5× MS agar plates. For fungal inoculation, a 5 mm plug from a 7-day-old PDA culture was placed at the centre of each plate and six sterile soybean seeds were arranged equidistantly around the plug, forming a circle. For exCFS treatments, 5 μL of exCFS dissolved in methanol (final concentration 0.5 mg mL^-1^) were spotted onto the plate surface, and six seeds were placed directly onto each spot. Plates containing only the sterilized seeds were used as control to assess the basal germination, while seeds deposited on 5 μL of methanol alone served as solvent controls. To assess stress mitigation, the same experimental setups were performed on 0.5× MS plates supplemented with CuSO₄ (1 mM and 3 mM) or NaCl (0.1 M and 0.25 M). Both concentrations were selected based on previous in vitro assay results, which demonstrated their toxicity to soybean seeds (data not shown). For each treatment and stress condition, plates containing only seeds were included as controls.

All assays were carried out in two independent experiments, each performed in triplicate. Plates were incubated in darkness at 25 °C for 7 days. After incubation, seed germination was evaluated by measuring primary root length and seedling fresh weight.

### Evaluation of *T. pseudofuniculosus* FRA effect on soybeans in pot experiments

Soybean seeds were surface-sterilized as described above and sown at a depth of 1 cm in autoclaved horticulture soil (140 g per pot) and irrigated regularly with deionized water to avoid water stress. Seeds were kept in darkness for three days to facilitate germination and subsequently grown in a greenhouse under natural light conditions. After germination, the non-germinated seeds were removed and only the uniformly developed seedlings were retained, resulting in a total of 24 plants per condition (one plant per pot). Twelve days after sowing, seedlings were inoculated with 10 mL of 1x PBS containing 10^6^ spores of *T. pseudofuniculosus* FRA, applied near the rhizospheric zone, or with 10 mL of exCFS dissolved in methanol at a concentration of 0.5 g L^-1^. Plants treated with 10 mL of 1× PBS or 10 mL of methanol served as solvent controls, while plants irrigated with water only were used as untreated controls. At 21 days after sowing, abiotic stress was imposed for 10 days. Copper stress was applied by irrigating plants with 10 mL of a 14 mM CuSO₄ solution, corresponding to a final concentration of 1 mM Cu²⁺ in the soil (34.96 mg CuSO₄·5H₂O per pot). In parallel, salt stress was induced by irrigating plants with 20 mL of 0.7 M NaCl, yielding a final soil concentration of 0.1 M NaCl (818 mg NaCl per pot). A schematic overview of the experimental design is shown in Figure 1. At the end of the treatment, plants were harvested and shoot length, root length, and fresh weight were recorded, and representative images were acquired. To verify fungal root colonization, 1.0 g of roots was surface-sterilized (70% v/v ethanol for 5 min, followed by rinsing with sterile water), homogenized in 5 mL of sterile 1× PBS, and plated on Rose Bengal agar (Smith and Dawson, 1944).

**Figure 1.**
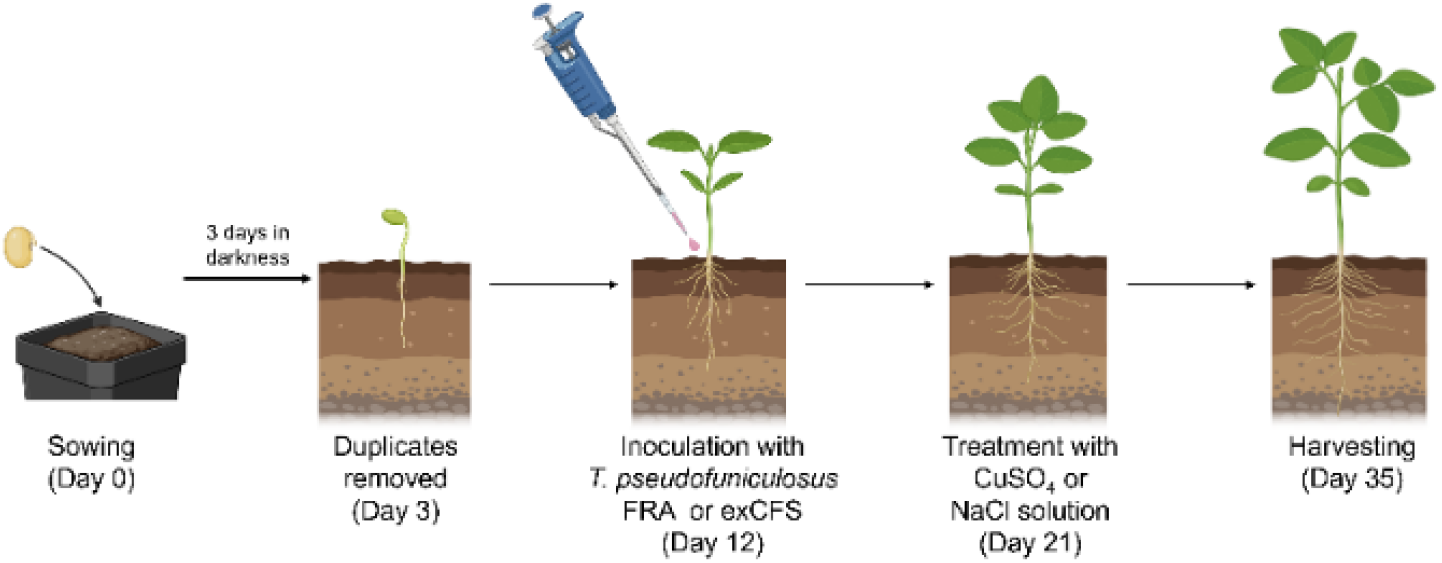
Schematic representation of pot experiments in greenhouse. Surface-sterile seeds were sown at 1 cm in autoclaved horticultural soil and kept in darkness for 3 days. After germination, only equally germinated seeds were kept in pots (1 seed per pot, 24 pots per condition). At day 12, plants were inoculated with a spore suspension of T. pseudofuniculosus FRA or treated with exCFS. At day 21 after sowing, plants were subjected to copper or salt stress. Plants were harvested 35 days after sowing.

### Statistical analysis

All statistical analyses were performed using GraphPad Prism software (version 8.0.1; GraphPad Software, San Diego, CA, USA), and data were expressed as mean ± standard error (S.E.). As indicated in the figure legends, differences among groups were analysed using one-way analysis of variance (ANOVA) followed by Tukey’s multiple comparison test or Dunnett’s test, or by a two-tailed unpaired Student’s t-test, as appropriate. A significance level of α = 0.05 was applied, and differences were considered statistically significant when p < 0.05.

## RESULTS

### Isolation and identification of *Talaromyces pseudofuniculosus* FRA

Samples of extremely acidic water (pH 2.13), collected from “Piccolo Inferno” (Campania, Italy), were used for fungal isolation to obtain extremophilic microorganisms capable of tolerating multiple abiotic stresses and potentially improving plant growth under adverse environmental conditions. After a 7-day enrichment at 25 °C in acidified water (pH 2.5), fungal growth was observed, indicating the presence of acidophilic heterotrophs. The isolate was transferred to PDA plates and incubated at 25 °C in darkness. Colony morphology analysis revealed velvety green, cottony colonies (Figure 2A), while microscopic observation (40×, Olympus-BH2) revealed branched conidiophores with abundant conidia (Figure 2B).

**Figure 2.**
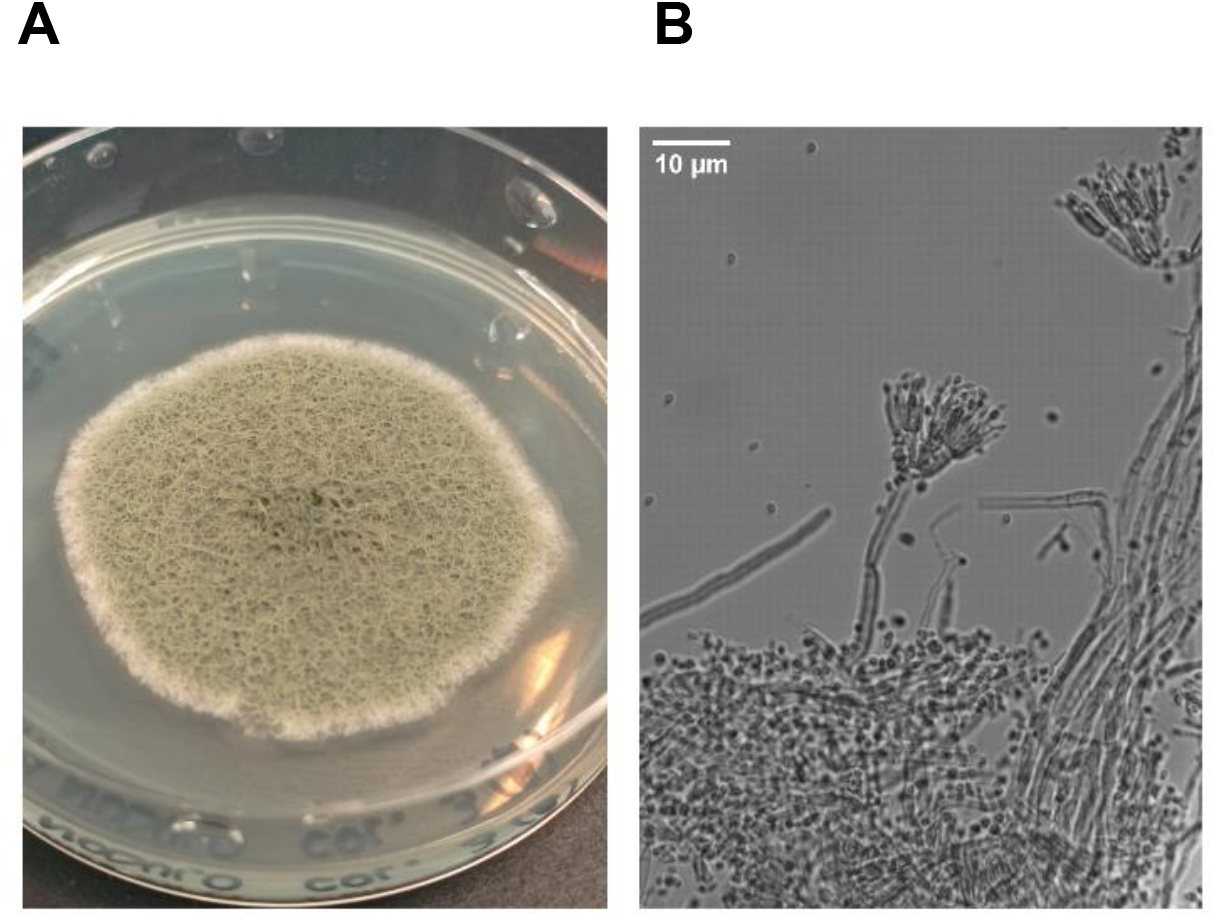
(A) Colony morphology of the fungal isolate on PDA after 7 days of growth at 25 °C. (B) Light microscopy image showing biverticillate conidiophores bearing numerous asexual conidia and septate hyphae (40× magnification).

Interestingly, the identification of the fungal isolate was performed using both the ITS region, the primary DNA barcode for fungal identification, and the β-tubulin (BenA) gene, a recommended secondary marker for species-level identification within the genus *Talaromyces* (Yilmaz *et al*., 2014). Following DNA extraction, both regions were amplified and sequenced, and a subsequent BLAST search was performed against the NCBI GenBank database. The ITS sequence analysis identified the fungus as *Talaromyces pseudofuniculosus* (formerly treated within *Penicillium* sensu lato (Yilmaz *et al*., 2015), exhibiting a 99.41% sequence identity with *Talaromyces pseudofuniculosus* FMR 15307. BenA sequence analysis further supported the species-level assignment, showing 99.32% sequence identity to the same *T. pseudofuniculosus* strain. Based on the combined molecular evidence, the isolate was designated as *Talaromyces pseudofuniculosus* FRA.

### *T. pseudofuniculosus* FRA is tolerant to multiple abiotic stressors

Fungi from extremely acidic environments often display cross-tolerance to several stresses due to shared physiological and genetic adaptations. Accordingly, the tolerance of *T. pseudofuniculosus* FRA to salinity and heavy metals was investigated by growing the fungus on PDA supplemented with NaCl (0.1–1.0 M) or increasing concentrations (1 -4 mM) of CuSO₄, PbCl₃, and NiSO₄. As shown in Figure 3A, *T. pseudofuniculosus* FRA presents a remarkable salt stress tolerance, evidenced by its ability to grow in the presence of up to 1M NaCl. Heavy metal tolerance assays revealed a metal-specific response (Figure 3B). At 1 mM, growth was comparable to the control in the presence of CuSO₄ and PbCl₃, whereas NiSO₄ caused a marked reduction in growth, with a tolerance index below 20%. Increasing NiSO₄ concentrations fully inhibited fungal growth. Copper caused a progressive reduction in growth, exceeding 80% inhibition at 4 mM, while lead resulted in approximately 50% inhibition at the same concentration.

**Figure 3.**
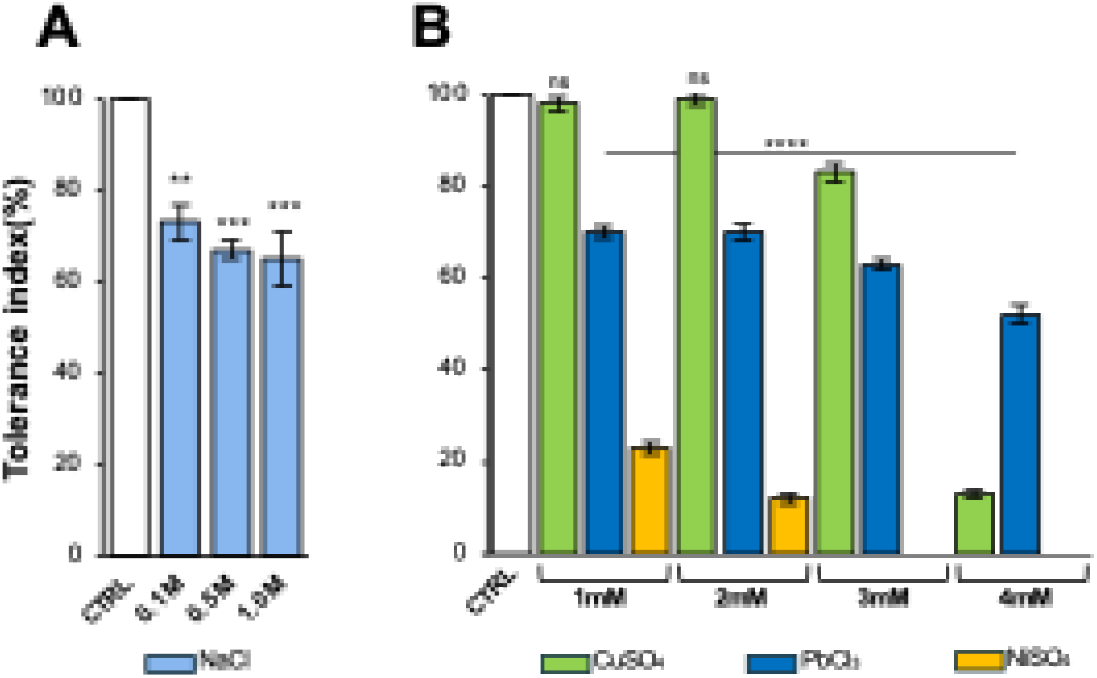
Effect of abiotic stress on radial growth of T. pseudofuniculosus FRA under different concentrations of NaCl (A) and heavy metals (B). Results are expressed as mean ± S.E. of three independent experiments. One-way ANOVA test was performed to compare the groups of data, and p-values are shown in the figure. Asterisks indicate significant differences in fungus growth compared to the respective control (CTRL, fungus grown without stress) according to Dunnet’s test; ns: not significant; *:: p < 0.0332; **: p < 0.0021; ***: p < 0.0002; ****: p < 0.0001.

In parallel, changes in culture pH during fungal growth under control conditions were monitored in PD broth at 7, 14, and 21 days. The initial pH of the medium was 5.9 and remained substantially unchanged after 7 days of growth. A marked acidification was observed at later stages, particularly between 14 and 21 days, with the culture reaching approximately pH 4 at day 21.

### *In vitro* evaluation of plant growth-promoting properties of *T. pseudofuniculosus* FRA

*T. pseudofuniculosus* FRA was evaluated for its PGP potential under stress conditions using salinity and Cu²⁺ as representative and agronomically relevant stressors.

#### Phosphate solubilization under salt and copper stress

Phosphate solubilization was evaluated on Pikovskaya medium supplemented with NaCl or CuSO₄. The phosphate solubilization index (P.S.I.) was calculated after 3 and 6 days of incubation. As shown in Figure 4A, increasing salinity did not reduce phosphate solubilization, and significantly higher P.S.I. values were observed at 1.0 M NaCl compared with the control at both time points (Figure 4A). Under copper stress, the P.S.I. at day 3 was slightly but significantly higher than that of the control (Figure 4B), indicating that copper at the tested concentrations did not impair phosphate solubilization. However, after 6 days of incubation, the P.S.I. decreased significantly under all copper concentrations tested.

**Figure 4.**
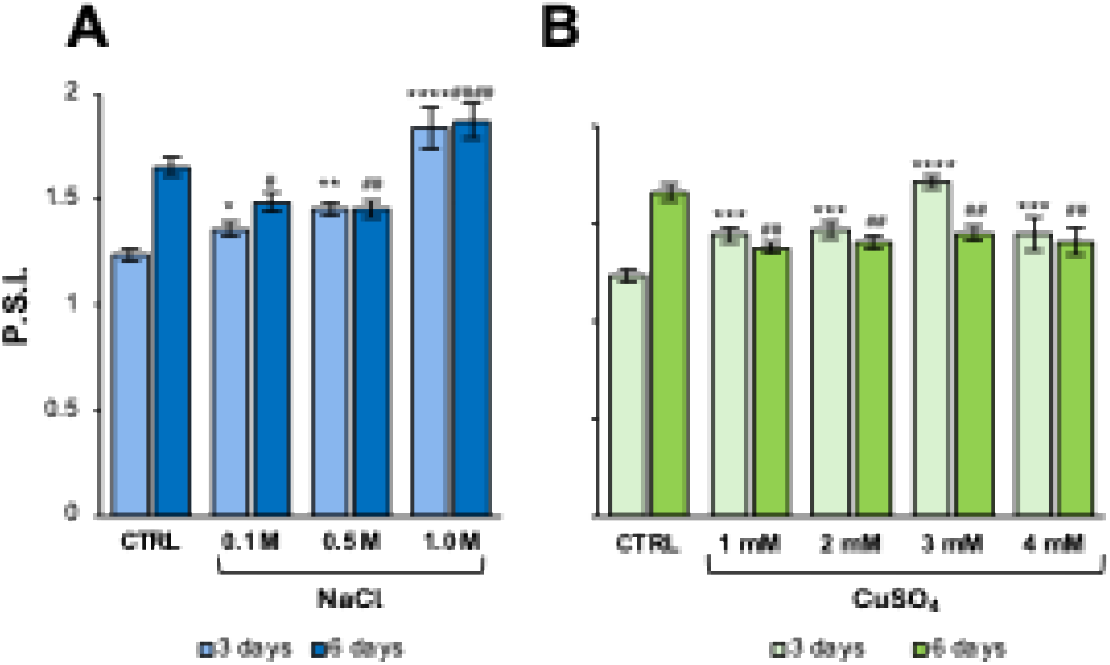
Phosphate Solubilization Index of T. pseudofuniculosus FRA under salt (A) and copper (B) stress. Bars represent the means ± S.E. of four independent experiments. Statistical differences were analysed by one-way ANOVA followed by Dunnett’s multiple comparison test against the respective control (CTRL). Asterisks indicate significant differences at 3 days, whereas hashes indicate significant differences at 6 days: ns: not significant; */ ^#^: p < 0.0332; **/ ^##^: p < 0.0021; ***/ ^###^: p < 0.0002; ****/ ^####^: p < 0.0001.

#### Production of Indole-3-Acetic Acid and IAA-Related Compounds under stress

Indole-3-acetic acid production, a key auxin mediating plant–microbe interactions, was quantified in Tryptophan Liquid Media under control, salt, and copper stress conditions. As shown in Figure 5, *T. pseudofuniculosus* FRA can produce IAA in TLM supplemented with Tryptophan even in the presence of the abiotic stresses. Moreover, the IAA production was clearly stress modulated, with the maximum production (423 mg L^-1^) obtained under low salt stress (0.1 M NaCl) after 14 days. Low amounts of CuSO_4_ (1 mM) allowed the production of a small amount of IAA after 14 days (90.6 mg L^-1^), while higher amounts of CuSO_4_ (3 mM) totally inhibited its production.

**Figure 5.**
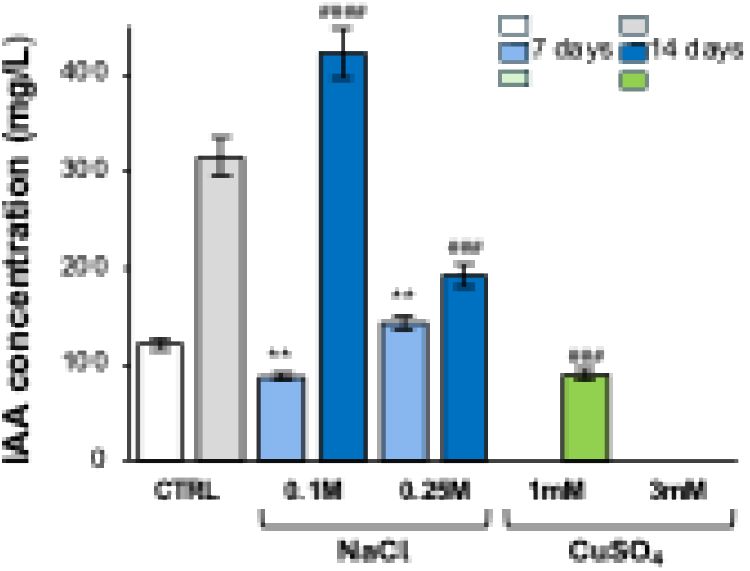
Indole-3-acetic acid production by T. pseudofuniculosus FRA grown in Tryptophan Liquid Medium under control conditions (CTRL), salt stress (NaCl 0.1 M and 0.25 M), and copper stress (CuSO₄ 1 mM and 3 mM). IAA concentrations were determined after 7 days (light bars) and 14 days (dark bars) of incubation and are expressed as mg/L. Bars represent the mean ± S.E. of three independent experiments. Statistical differences were analysed by one-way ANOVA followed by Dunnett’s multiple comparison test against the respective control (CTRL). Asterisks indicate significant differences at 7 days, whereas hashes indicate significant differences at 14 days: ns: not significant; */ ^#^: p < 0.0332; **/ ^##^: p < 0.0021; ***/ ^###^: p < 0.0002; ****/ ^####^: p < 0.0001.

### *T. pseudofuniculosus* FRA hydrolytic and biocontrol activities

To characterize the hydrolytic-enzymatic capabilities of the isolate, *T. pseudofuniculosus* FRA was tested for cellulase, amylase, and xylanase activities under control conditions and under salt and copper stress. To this aim, carboxymethyl-cellulose (MMCMC) media, soluble starch media (MMAMY), and xylan media (MMXYL) were used to evaluate cellulase, amylase, and xylanase activity, respectively. After a 7-day incubation period, hydrolytic activities were quantified as the Enzymatic Index (EI), as described in the Materials and Methods section. The appearance of clear hydrolysis halos on MMCMC and MMAMY plates confirmed the ability of *T. pseudofuniculosus* FRA to degrade cellulose and starch, with EI values of 1.89 and 1.82, respectively, whereas no xylanase activity was detected. Under abiotic stress conditions, amylase activity was completely inhibited by both NaCl and CuSO₄. In contrast, cellulolytic activity was maintained under saline stress, although progressively reduced with increasing NaCl concentrations, whereas copper completely inhibited cellulose hydrolysis (Figure 6).

**Figure 6.**
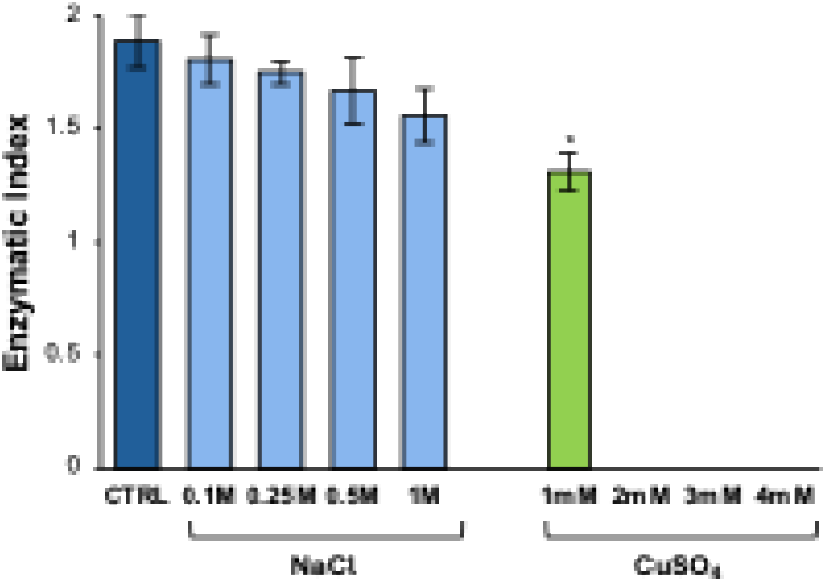
Cellulase activities of T. pseudofuniculosus FRA expressed as Enzymatic Index. Cellulase activity evaluated under control conditions (CTRL) and under salt stress (NaCl, 0.1, 0.25, 0.5, and 1.0 M) and copper stress (CuSO₄, 1, 2, 3, and 4 mM) after 7 days of incubation at 25 °C. Bars are reported as mean ± S.E. of four independent experiments. Statistical differences were analysed by one-way ANOVA followed by Dunnett’s multiple comparison test against the respective control. *: p < 0.0332.

Since *T. pseudofuniculosus* FRA retained cellulolytic activity under salt stress, its endo- and exoglucanase activities were quantified as described by Meng et al. (2021). Quantitative assays revealed high glucanase production, with endoglucanase and exoglucanase activities of 14 ± 0.14 U mL⁻¹ and 7 ± 0.1 U mL⁻¹, respectively.

Since hydrolytic activities can be associated with antifungal activity (Panchal, 2022; Ragucci *et al.,* 2023), the biocontrol potential of the fungus was evaluated to determine whether it could also confer protection against biotic stresses caused by phytopathogen fungi (Supplementary materials). Although no direct antifungal activity was detected by dual culture assays (Figure S1), competitive overgrowth suggested a potential indirect biocontrol capacity.

### Extracellular metabolite profiling of *T. pseudofuniculosus* FRA

The secreted secondary metabolites of *T. pseudofuniculosus* FRA were identified through a gas chromatography coupled with mass spectroscopy analysis (GC-MS). The secreted molecules were recovered from the CFS after 7 days of fungal growth and analysed by GC–MS after trimethylsilylation, as described in the Materials and Methods. The total ion current chromatogram of the exCFS is shown in Figure 7. Sixteen compounds were identified (Table S1), including several organic acids and their derivatives, such as lactic acid, β-lactic acid, 3-hydroxyisobutyric acid, pyroglutamic acid, 5-hydroxymethyl-2-furoic acid, 2-isopropylmalic acid, 2-hexylbutanedioic acid, citric acid, and gluconolactone. Other detected compounds included glycerol, phenylacetic acid, uracil, furandimethanol, orcinol, 4-hydroxybenzoic acid, and 2′,4′,6′-trihydroxyacetophenone.

**Figure 7.**
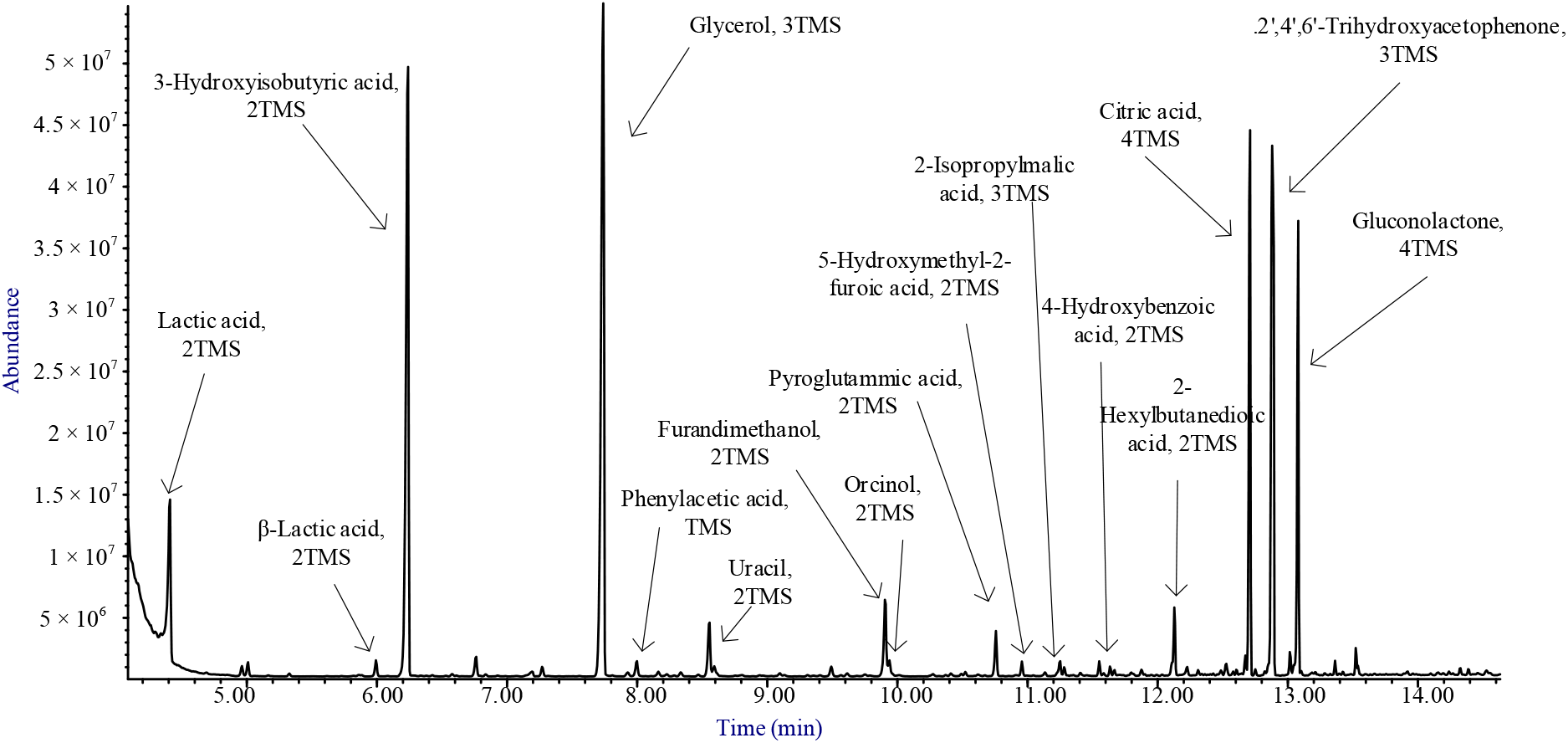
Total ion current chromatogram of crude extract of T. pseudofuniculosus FRA after derivatization with BSTFA.

### *T. pseudofuniculosus* FRA and exCFS display *in vitro* plant growth promotion properties on *Glycine max* L

To confirm the beneficial effects of *T. pseudofuniculosus* FRA observed *in vitro*, the potential PGP activity of the live fungus and its cell-free supernatant extracts were evaluated using Soybean (*Glycine max* L.) as a model plant system. Their plant growth-promoting and stress-mitigating potential on early seedling development was assessed under control conditions and in the presence of NaCl or CuSO₄ stress. After seven days, seedling fresh weight and primary root length were measured as indicators of growth performance (Figure 8).

**Figure 8.**
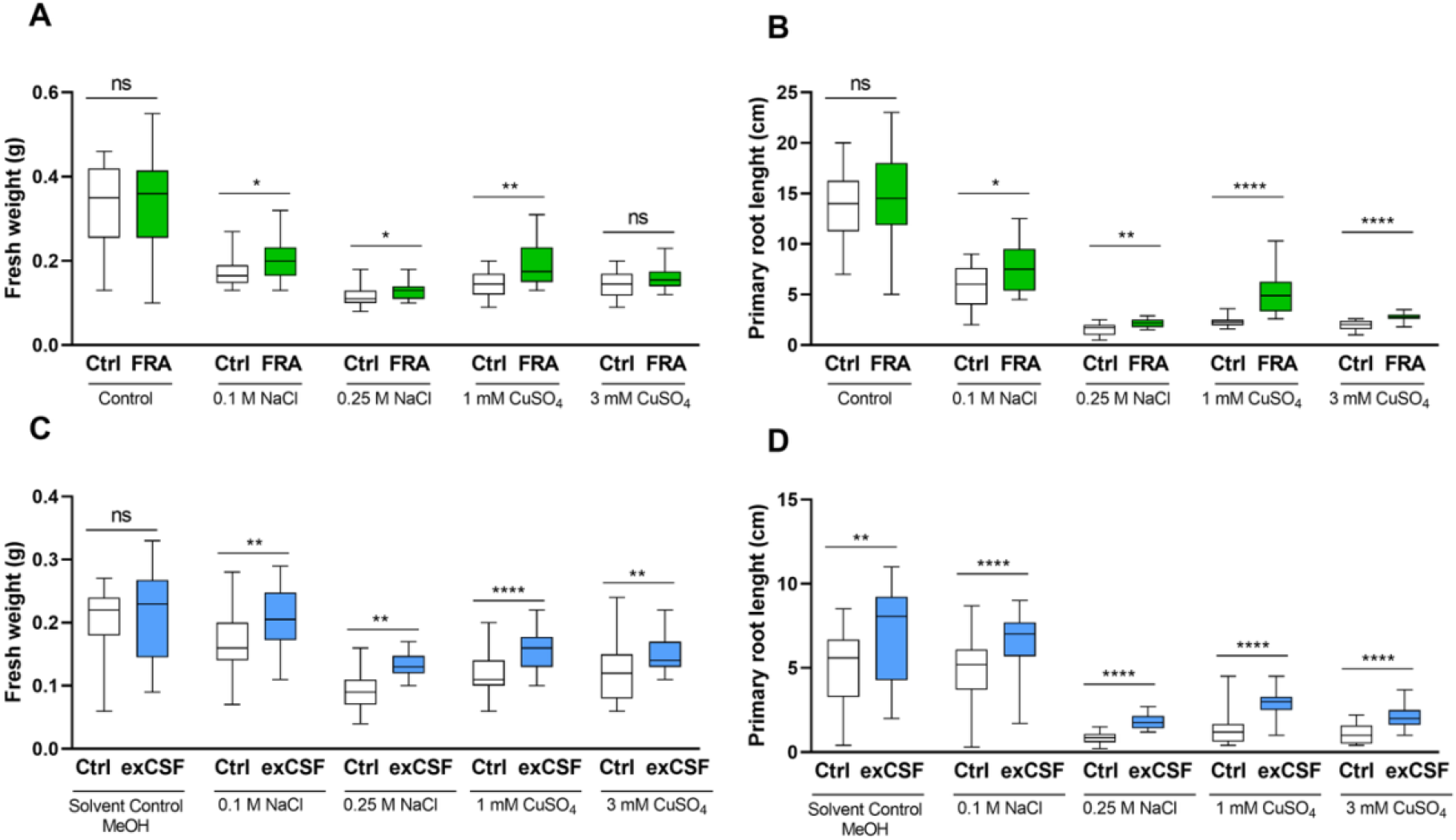
Effect of T. pseudofuniculosus FRA and its cell-free supernatant extract on Glycine max L. seedling growth under abiotic stress. Seedlings were treated with the live fungus (FRA)(A, B) or exCFS (C, D) and seedling fresh weight (A, C) and primary root length (B, D) were measured after 7 days under control, NaCl (0.1 and 0.25 M), or CuSO₄ (1 and 3 mM) conditions. Boxes indicate the interquartile range (25^th^-75^th^ percentile) with the median (line) and whiskers indicate the minimum and maximum values. n = 24 (four independent experiments, 6 seeds per plate). Statistical analysis was performed using unpaired t-test, comparing each treatment with fungus or exCFS (+) with the respective control (-). Statistical significance is reported with asterisks on the graphs, ns: not significant; *: p < 0.0332; **: p < 0.0021; ***: p < 0.0002; ****: p <0.0001.

As reported in Figure 8A-B, in the absence of stress, the fungal inoculation (green boxes) did not significantly affect either parameter compared to the non-inoculated seeds (white boxes). Under salinity stress, however, the presence of the fungus slightly but significantly enhanced seed growth in terms of both weight and length (Fig. 8A-B). Copper stress strongly reduced seedling development, which was partially restored by the fungus. In particular, under moderate copper stress (1 mM CuSO_4_), fungal inoculation significantly improved both seedling length and weight. At higher copper stress (3 mM CuSO_4_), fungal inoculation no longer affected seedling biomass but continued to promote root elongation, suggesting a residual growth-stimulatory effect even under severe stress conditions. Figure 8C-D shows the impact of exCFS (dissolved in MeOH at a concentration of 0.5 mg mL^-1^) on seedling biomass (C) and primary root length (D). When the seedlings were treated with exCFS, under control conditions (control with MeOH), exCFS inoculation (blue boxes) did not significantly alter seedling weight but significantly increased seedling length. Under all tested abiotic stress conditions, exCFS treatment significantly improved both the length and the biomass compared with the corresponding controls. Overall, both fungal inoculation and exCFS application enhanced soybean seedling growth under abiotic stresses. Although general growth values were lower in the exCFS assay due to methanol, exCFS-treated seedlings consistently performed better than their respective solvent controls, confirming the positive biological activity of the fungal metabolites. Considering these results, both the fungal supernatant extract and the fungus itself exhibit promising potential as biostimulants for *Glycine max* L. in the presence of salt or copper contamination.

### Evaluation of *T. pseudofuniculosus* FRA and exCFS as Growth Promoters in Soybean Pot Trials

After *in vitro* tests, the positive effects of both *T. pseudofuniculosus* FRA and exCFS were evaluated on soybean plant growth under salt and copper stress in pot experiments. These experiments aimed to confirm whether the plant growth-promoting effects observed *in vitro* could be reproduced under soil conditions, where complex microbe–plant and metabolite–root interactions take place. Plants were inoculated with either a spore suspension of *T. pseudofuniculosus* FRA or treated with its exCFS and subsequently exposed to salt (0.1 M NaCl) or copper (1 mM CuSO₄) stress conditions. After treatment, plants were harvested, and morphological parameters were evaluated.

The association of *T. pseudofuniculosus* FRA with soybean plants significantly increased plant weight, leaf area, and plant height under all the conditions tested, as reported in Figure 9, compared to the non-inoculated plants. In the absence of stress,

**Figure 9.**
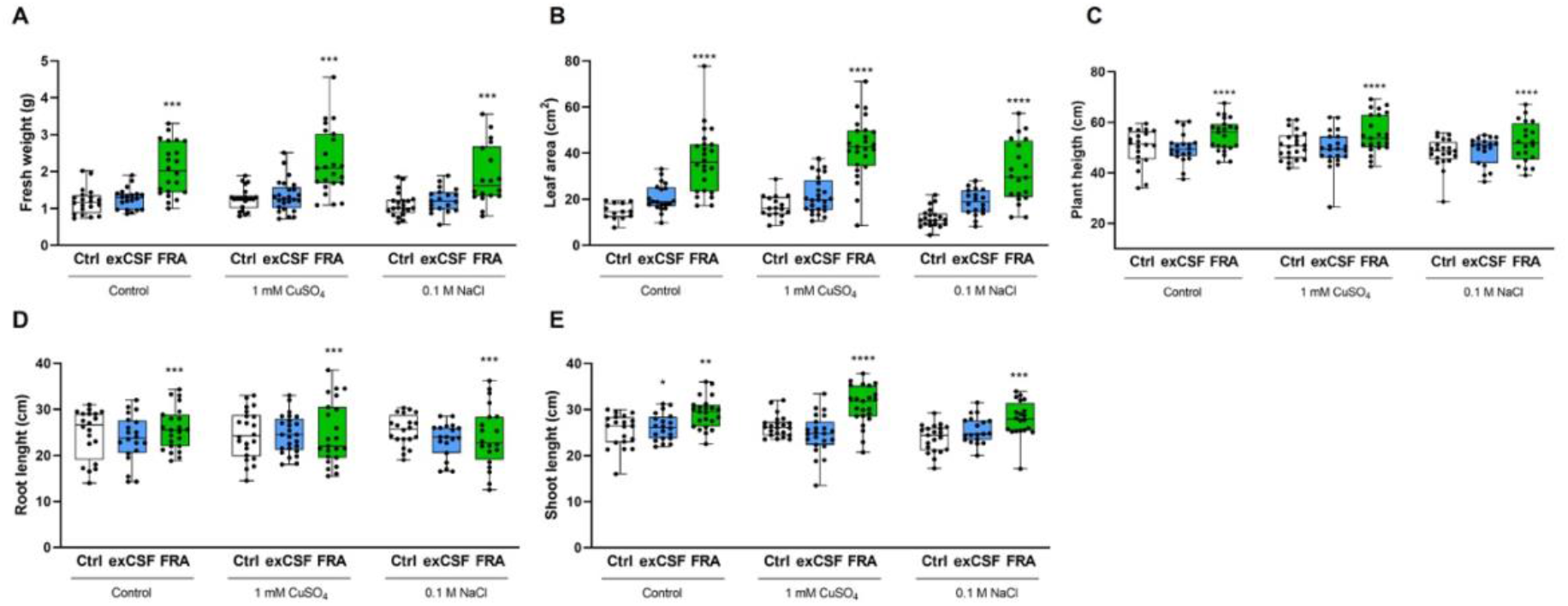
Effect of T. pseudofuniculosus FRA on soybean growth under abiotic stress in pot experiments. Plant fresh weight (A), leaf area (B), and plant height (C) were measured in soybean plants treated with the live fungus (FRA) or its cell-free supernatant extract under control, copper (1 mM CuSO₄), and salt (0.1 M NaCl) conditions. Plant height was further separated into root (D) and shoot (E) length. Results are reported as mean (n = 24) ± S.E. of independent experiments (one plant per pot). One-way ANOVA test was performed to compare the groups of data, and p-values are shown in the figure. Asterisks indicate significant differences of FRA and exCFS with the respective control according to Tukey’s test, ns: not significant; *: p < 0.0332; **: p < 0.0021; ***: p < 0.0002; ****: p < 0.0001.

*T. pseudofuniculosus* FRA inoculation resulted in a substantial enhancement of 74% in plant weight, 149% in leaf area, and 20% in plant height, in comparison to non-inoculated plants. Under copper stress, *T. pseudofuniculosus* FRA enabled soybean plants to withstand metal-induced toxicity, leading to increases of 78% in plant weight, 153% in leaf area, and 17% in plant height. Similarly, under salt stress, inoculation improved plant growth by 70% for plant weight, 166% for leaf area, and 20% for shoot length. In contrast, application of exCFS did not result in significant improvements in whole-plant morphology in pot experiments.

### Root colonization and qualitative GC–MS profiling under stress

To investigate the different responses observed in plants under stress inoculated with *T. pseudofuniculosus* FRA spores or treated with exCFS, root colonization and extracellular metabolite profiles were assessed. Root colonization analysis showed that the FRA strain was able to establish within soybean roots, as the fungus was recovered exclusively from roots of inoculated plants (Figure S2). In parallel, extracellular metabolites produced under NaCl (0.1 M) and CuSO₄ (1 mM) conditions were analysed by GC–MS after 7 days of fungal growth (Fig. S3). The results suggest that *T. pseudofuniculosus* FRA adjusts its metabolism according to the growth environment. However, comprehensive metabolomic studies are required to quantify the produced metabolites.

## DISCUSSION

In this study, we isolated a fungal strain from the acidic waters of Piccolo Inferno (Italy) and identified it as *Talaromyces pseudofuniculosus* FRA based on ITS and β-tubulin sequence analyses. The isolate exhibited multi-stress tolerance together with several plant growth-promoting traits, which enabled the strain to enhance soybean tolerance to salinity and copper stress. Previous studies have reported similar features in closely related *Talaromyces* species, such as *T. funiculosus* (Khan et al., 2011; Khan and Lee, 2013; Bilal et al., 2019). However, to the best of our knowledge, this is the first report describing in *T. pseudofuniculosus* a multi-stress-tolerant phenotype combining tolerance to salinity and different heavy metals with the maintenance of plant growth-promoting activities under stress conditions.

Fungi inhabiting extremely acidic environments frequently exhibit cross-resistance to multiple environmental stressors as a consequence of shared adaptive mechanisms related to osmotic balance, antioxidant responses and metal detoxification (Das and Roychoudhury, 2014; Das *et al*., 2022). The origin of *T. pseudofuniculosus* FRA from an environment with a pH of 2.13 is therefore consistent with the pronounced tolerance to salinity and metals observed in this study (Fig. 3A-B). The different responses to Ni²⁺, Cu²⁺ and Pb²⁺ further indicate that this tolerance is metal-specific rather than a generalized resistance phenotype. Importantly, despite originating from an extremely acidic habitat, FRA was also able to grow under non-stress conditions and remained physiologically active under salinity and copper stress. Such ability to tolerate extreme conditions while retaining functionality under more favorable environments has been recognized as a strategic advantage of extremophilic and extremotolerant fungi for agricultural applications (Yarzábal Rodríguez et al., 2024).

Among the PGP traits displayed by FRA, phosphate solubilization, phytohormone production and hydrolytic activities provide complementary mechanisms through which the fungus could support plant growth. *T. pseudofuniculosus* FRA retained its ability to solubilize inorganic phosphate under salinity and moderate copper stress, indicating that nutrient mobilization remains functional under environmental stress (Fig. 4 A-B). Soil fungal species, including species of *Penicillium*, *Aspergillus*, *Rhizopus* and *Fusarium*, commonly mobilize poorly available phosphate through secretion of low-molecular-weight organic acids (Whitelaw, 1999; Rashid *et al*., 2004), and several such compounds were detected in the FRA extracellular extract by GC–MS (Fig. 7, Table S1). These compounds, such as citric, lactic, acetic and succinic acids, may contribute to phosphate mobilization through acidification and chelation and may also affect nutrient availability in the surrounding environment, influencing endemic microbiota (Vassileva et al., 2022). The accumulation of organic acids in filamentous fungi is strongly influenced by the availability of metal ions, including Cu²⁺, which can affect fungal physiology and primary metabolism (Karaffa et al., 2021). Accordingly, the decrease in phosphate solubilization observed after prolonged copper exposure may reflect a stress-dependent modulation of fungal metabolism and, potentially, of organic acid production (Antsotegi-Uskola et al., 2020; Karaffa et al., 2021). IAA production was also strongly dependent on incubation time and environmental conditions (Fig. 5). IAA is a key auxin mediating plant–microbe interactions, able to modulate root architecture and nutrient acquisition (Arora *et al*., 2024). Notably, *T. pseudofuniculosus* FRA maintained auxin-related production under both salinity and moderate copper stress. The highest production was observed under 0.1 M NaCl after 14 days, reaching 423 mg L⁻¹, whereas substantial production was still detected at 1 mM CuSO₄ (90.6 mg L⁻¹), while 3 mM CuSO₄ completely inhibited it. Stress-dependent IAA production has also been reported in plant-beneficial fungi: *Trichoderma harzianum* InaCC F88 maintained IAA production under increasing NaCl concentrations (Napitupulu *et al*., 2019), while IAA production under Cu exposure has been demonstrated in the culture filtrates of *Porostereum spadiceum* AGH786 and the Cu-tolerant endophyte *Aspergillus tamari* SK12 (Naz et al., 2022; Khan et al., 2024). Although differences in culture media, tryptophan availability, incubation time and analytical conditions limit direct quantitative comparisons among studies, the levels detected in the FRA strain under stress conditions compare favorably with those reported for these fungi. These results highlight the strong auxin-producing capacity of FRA and its ability to maintain this PGP function under distinct abiotic stresses. In addition, the FRA strain also showed cellulolytic and amylolytic activities, with cellulase activity being retained even at high NaCl concentrations (Fig. 6). Extracellular hydrolytic enzymes produced by filamentous fungi contribute to organic matter decomposition and nutrient turnover, and their maintenance under stress may further increase nutrient availability in marginal soils (Ogunmolu et al., 2015).

GC–MS analysis of the exCFS provided additional insights into the extracellular chemistry potentially underlying these activities. The extract contained organic acids, polyols and aromatic compounds with possible roles in nutrient mobilization, metal complexation and stress responses (Table S1). Lactic acid, β-lactic acid, 3-hydroxyisobutyric acid, citric acid and gluconolactone are consistent with mechanisms involving rhizosphere acidification and mineral nutrient mobilization, whereas citric acid and related organic acids may also contribute to metal complexation. Glycerol may participate in osmotic stress protection, while phenylacetic acid, an auxin-related compound, may influence root development. Rather than indicating a single mechanism, the extracellular profile therefore supports a multifactorial contribution of fungal metabolism to nutrient availability and plant stress responses.

The plant assays confirmed the functional relevance of these PGP traits (Fig. 8, 9). During early soybean development, the live fungus had limited effects under non-stress conditions but significantly enhanced seedling growth under salinity and copper stress (Fig. 8 A-B), indicating that its beneficial effects become particularly evident under environmental challenge. The exCFS also improved seedling length and biomass under the tested stress conditions (Fig. 8 C-D). Although absolute growth values were lower in these experiments, likely because of the methanol used for extract solubilization, exCFS-treated seedlings consistently outperformed their corresponding solvent controls. Thus, secreted fungal metabolites alone are sufficient to exert measurable biological activity during germination.

A different pattern emerged in pot experiments (Fig. 9). Fungal inoculum produced sustained increases in soybean fresh weight, leaf area and plant height under both control and stress conditions (Fig. 9 A-C), with the response being mainly associated with shoot rather than root elongation (Fig. 9 D-E). In contrast, the positive effects of exCFS previously observed were not maintained at the whole-plant level in pot experiments (Fig. 8 C-D, 9). This difference suggests that a single application of extracellular metabolites may influence early developmental processes but but cannot fully reproduce the sustained effects of a metabolically active fungal inoculum.

The importance of an active fungus–plant association is further supported by the recovery of FRA from inoculated soybean roots, demonstrating successful root colonization. Persistent colonization has also been associated with plant growth stimulation by beneficial endophytic fungi such as *Trichoderma* (Csótó et al., 2023), suggesting that establishment within plant tissues may contribute to prolonged beneficial effects. In addition, exposure to NaCl and CuSO₄ indicates that the extracellular metabolism of FRA is dynamically modulated by environmental conditions, consistent with the recognized metabolic plasticity of filamentous fungi (Lin et al., 2024).

Together, the comparison between live FRA and exCFS highlights a potentially important feature of microbial inoculants. In contrast to a cell-free preparation, which contains a fixed set of metabolites generated under defined culture conditions, a living fungal inoculum can continuously respond to environmental and plant-derived signals and adjust its metabolism accordingly. The combination of root colonization, maintenance of PGP traits under stress, and stress-dependent changes in the extracellular metabolite profile therefore supports the hypothesis that physiological and metabolic plasticity contributes to the sustained plant growth-promoting effect of FRA. Nevertheless, the GC–MS analysis performed here provides a preliminary qualitative analysis of extracellular metabolite profiles rather than a quantitative metabolomic analysis. Targeted quantitative metabolomics, together with functional assays of selected metabolites and analyses directly within the plant–fungus system, will be required to establish causal relationships between specific fungal metabolites and the observed PGP effects.

## CONCLUSIONS

In this study, *Talaromyces pseudofuniculosus* FRA, isolated from the extremely acidic “Piccolo Inferno” environment, emerged as a physiologically and metabolically versatile plant growth-promoting fungus. Despite its extreme origin, FRA remained functional under non-stress conditions and tolerated distinct abiotic stresses, including salinity and heavy metals, while maintaining key PGP activities. Its ability to mobilize nutrients, produce auxin-related compounds and hydrolytic enzymes, colonize soybean roots, and modulate its extracellular metabolite profile in response to NaCl and CuSO₄ highlights the adaptive plasticity of the strain. Importantly, the more sustained effects of the live fungus compared with exCFS emphasize the relevance of an active and dynamic plant–fungus association for long-term growth promotion under stress. Overall, these findings support *T. pseudofuniculosus* FRA as a promising candidate for the development of fungal biostimulants for saline- and metal-impacted agricultural soils.

## Notes

### Competing Interest Statement

The authors have declared no competing interest.

### Funding

- Program for the Finanziamento della Ricerca di Ateneo (FRA) 2022 52 dell’Università degli Studi di Napoli Federico II.

