## supplemental materials for "From acidic extreme environments to agriculture: a multi-stress-tolerant *Talaromyces pseudofuniculosus* enhances soybean growth under abiotic stress"

**SUPPLEMENTARY MATERIALS**

### Biocontrol activity of *T. pseudofuniculosus* FRA

The antifungal activity of *T. pseudofuniculosus* FRA was evaluated using the dual-culture plate method against the phytopathogenic fungi *Alternaria alternata* (Castaldi *et al.*, 2023), *Macrophomina phaseolina* (Masi *et al.*, 2021), *Cercospora nicotianae* (Sautua *et al.*, 2020), *Colletotricum truncatum* (Ramos *et al.*, 2013), and *Septoria nodorum* (Ragucci *et al.*, 2023). These fungi were used because they are part of the collection held by the laboratory of Rachele Isticato at the Department of Biology, University of Naples “Federico II”. In brief, a 5 mm fungal disc of *T. pseudofuniculosus* FRA was deposited from 1.5 cm to each phytopathogenic fungal disc (5 mm) on PDA plates (ø 5 cm). As a control, the fungal plug from each fungus was positioned at the centre of each plate. All the plates were incubated at 25 °C for 10 days (Rahman *et al.*, 2016); after incubation, the growth of the two fungi was evaluated on each plate.

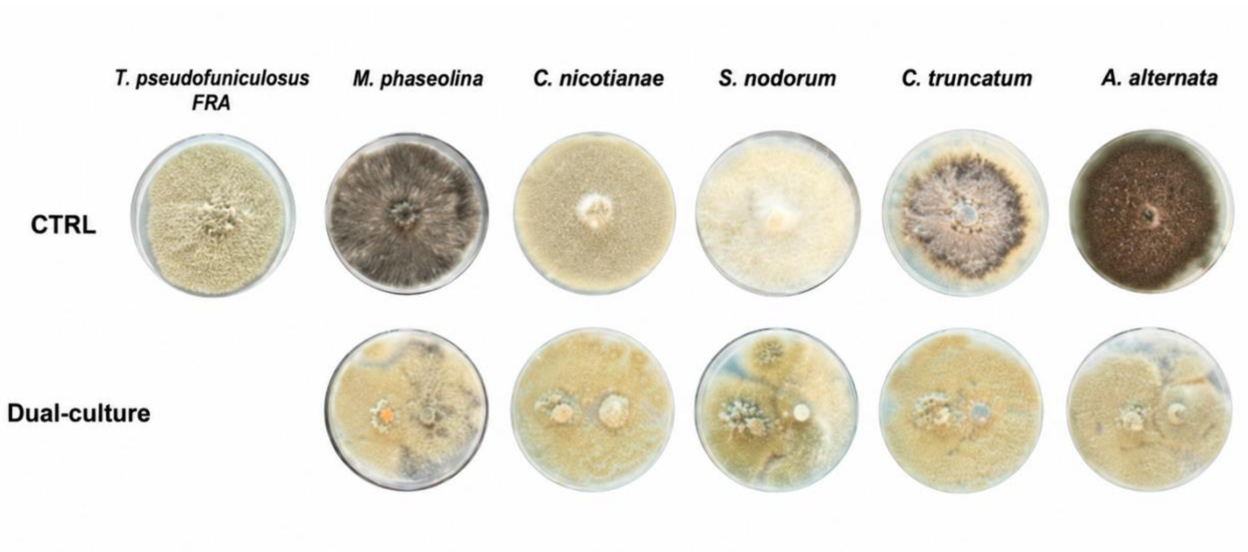


***Figure S 1.*** *Antifungal activity of T. pseudofuniculosus FRA against the phytopathogenic fungi M. phaseolina, C. nicotianae, S. nodorum, C. truncatum, A. alternata.*

**Table S1. Metabolites identified via GC-MS in exCFS of *T. pseudofuniculosus* FRA and their potential biological role**

| Compound | Chemical Class | Reported/Putative Role in Plant–Soil System |
|---|---|---|
| Lactic acid | Organic acid | Contributes to medium acidification; may modulate soil microbiome composition and act as carbon source for beneficial acidophilic bacteria (Vassileva et al., 2022). |
| β-Lactic acid | Organic acid | Selective nematicidal activity against plant-parasitic nematodes (Schwarz et al., 2004). |
| 3-Hydroxyisobutyric acid | Organic acid | May be associated with metabolic adaptation under energy/carbon stress (Schertl,et al., 2017). |
| Glycerol | Polyol | Enhances plant tolerance to abiotic and biotic stresses; reported to induce resistance against phytopathogens (Śliżewska et al., 2022). |
| Phenylacetic acid | Auxin-like Phytohormone | Promotes seed germination and early plant growth; modulates root development (Perez et al., 2023). |
| Uracil | Pyrimidine base | Involved in plant nucleotide metabolism and potentially in rhizosphere metabolic exchange (Girke et al., 2014,). |
| 2,5 Furandimethanol | Furan derivative | Potential antimicrobial activity; may indirectly shape soil microbiota (Dai et al., 2020). |
| Orcinol | Phenolic compound | Known antimicrobial properties; may contribute to indirect biocontrol activity (Nguyen et al., 2022). |
| Pyroglutamic acid | Organic acid derivative | May act in stress-response pathways and nitrogen metabolism (Lei et al., 2024). |
| 5-Hydroxymethyl--2-furoic acid | Furan derivative | Possible antioxidant and antimicrobial activity; may enhance host resistance (Chacar et al., 2025). |
| 2-Isopropylmalic acid | Phenolic acid | Involved in amino acid biosynthesis; may contribute to metabolic interactions in soil (Vigueras-Meneses et al., 2022). |
| 4-Hydroxybenzoic acid | Phenolic acid | Involved in modulating soil microbial community structure and displaying antibacterial/anti-virulence activity against plant pathogens (Guo et al., 2015; Kang et al., 2020). |
| 2-Hexylbutanedioic acid | Dicarboxylic acid | Its biological function in plant–soil systems remains poorly characterized. |
| Citric acid | Organic acid | Enhances plant growth and photosynthesis; promotes |

| | | |
|---|---|---|
| | | osmoregulation; chelates heavy metals, improving tolerance and soil detoxification (Tahjib-Ul-Arif et al., 2021). |
| 2',4',6'-Trihydroxy acetophenone | Phenolic compound | reported phytotoxic and seed-germination inhibitory activity (Chung et al., 2008). |
| Gluconolactone | organic acid | Contributes to acidification; may influence mineral Solubilization derivative and microbial community shifts (Suarez et al., 2019) |

**Verification of root colonization by *T. pseudofuniculosus* FRA**

Root colonization by *Talaromyces pseudofuniculosus* FRA was confirmed by re-isolation from surface-sterilized root tissues plated on Rose Bengal agar (Smith and Dawson, 1944). Abundant fungal growth was consistently recovered from roots of inoculated plants under control as well as NaCl and $CuSO_4$ treatments, whereas no colonies were detected in none of Mock or exCFS samples, confirming the effectiveness of the surface sterilization and excluding external contamination (Figure S2). These results demonstrate that *T. pseudofuniculosus* FRA is able to enter and persist within root tissues even under abiotic stress conditions, supporting a stable plant–fungus interaction underlying the growth-promoting effects observed in pot experiments

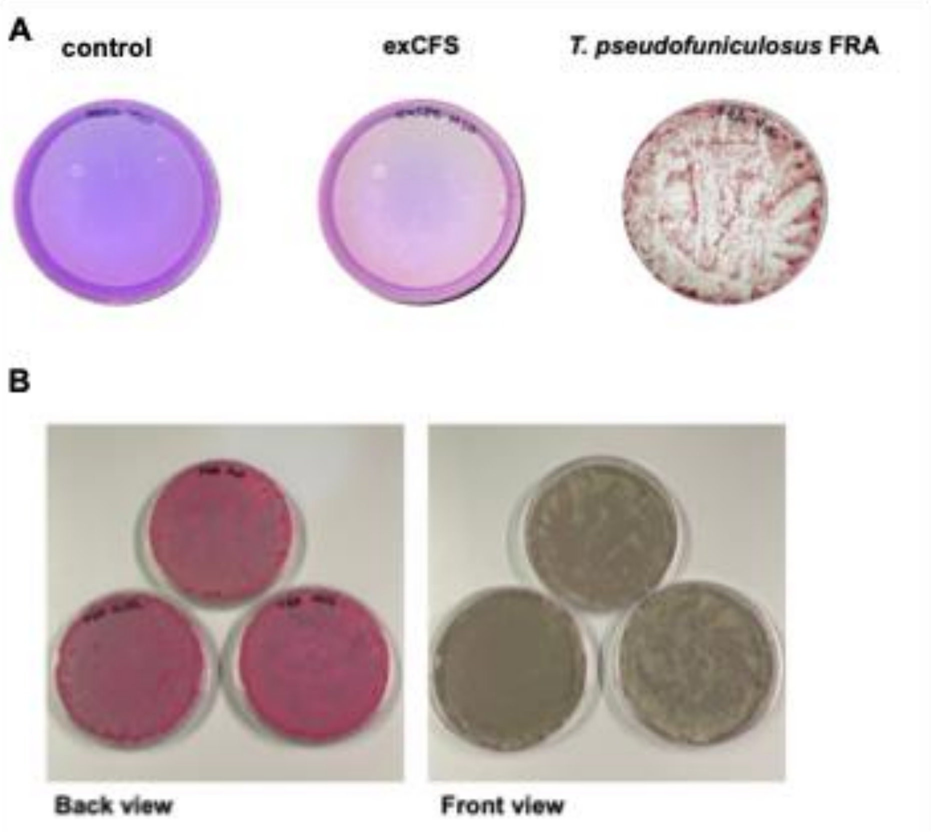


***Figure S 2.*** *(A) Re-isolation from surface-sterilized roots of control-, exCFS-, and FRA-treated plants grown under control conditions. (B) Re-isolation from roots of T. pseudofuniculosus FRA-inoculated plants grown under Mock ($H_2O$), NaCl, and $CuSO_4$ treatments. Roots (1 g) were surface sterilized, homogenized, and plated on Rose Bengal agar (Smith and Dawson, 1944). Fungal colonies were recovered exclusively from plants inoculated with the live strain, confirming internal colonization under both non-stress and stress conditions.*

### Extracellular metabolite profiling under abiotic stress conditions

A)

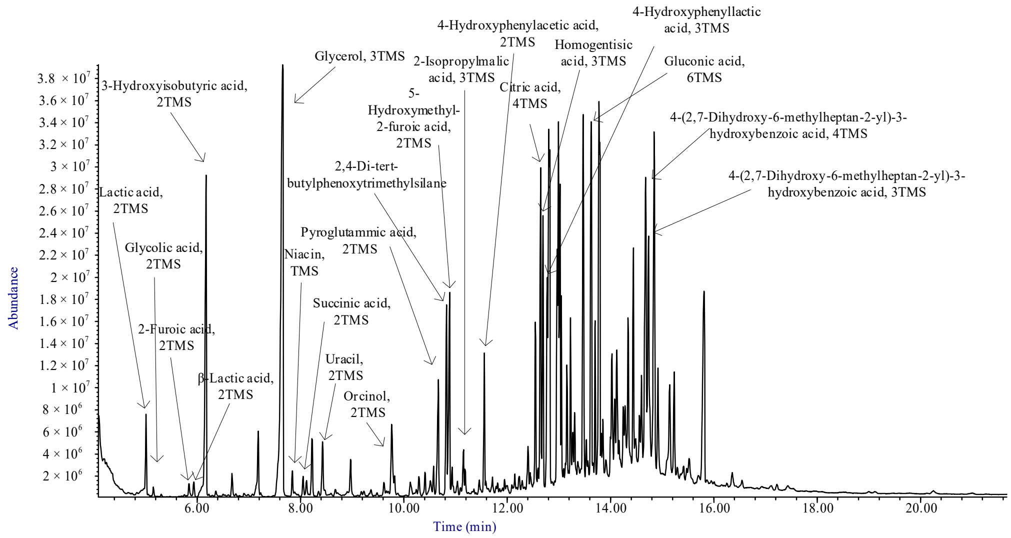


B)

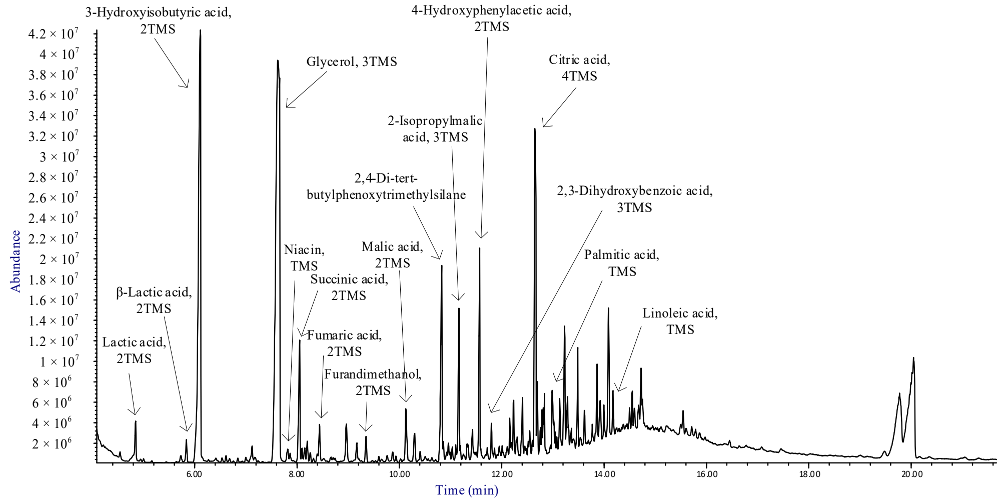


***Figure S3****. Total ion current chromatograms (TICs) of culture filtrate extracts of T. pseudofunuculosus FRA grown in presence of 0.1M NaCl (A) and 1mM $CuSO_4$ (B). GC-MS analyses were conducted after trimethylsilylation with BSTFA.*
